# Vicious cycle of hemodynamic perturbation and endothelial injury in development and progression of pulmonary arterial hypertension

**DOI:** 10.1101/2023.10.16.562339

**Authors:** Yupu Deng, Ketul R. Chaudhary, Anli Yang, Kirishani Kesavan, Liyuan Wang, Kevin Chathely, Duncan J. Stewart

## Abstract

**Background:** Pulmonary arterial hypertension (PAH) is a devastating disease caused by loss of effective lung microvasculature for which there is no curative treatment. Evidence from preclinical models and human disease-causing genetic mutations point to endothelial cell (EC) injury and apoptosis as a central trigger for the initiation of PAH. However, how EC apoptosis leads to pulmonary hypertension (PH) and complex arteriolar remodeling is uncertain.

**Methods:** Rats were subjected to SU5416-hypoxia (SUHx) and EC apoptosis, pulmonary vascular remodeling and arterial volume was assessed by immunohistochemistry, histology and microCT, respectively. Left pulmonary artery banding (LPAB) was performed, either 1 week before (prevention) or 5 weeks after SU injection (reversal), to study the effect of hemodynamic offloading.

**Results:** In the SUHx model, EC apoptosis was markedly increased as early as 3 days post-SU, persisting through PAH development, and this was associated with a profound arterial pruning with reduction in lung arterial volume (∼80%). LPAB abrogated lung EC apoptosis in the banded left lung and prevented as well as reversed arteriolar pruning. Moreover, in the reversal protocol, removal of the band at 10 weeks resulted in improvement in pulmonary hemodynamics and RV function at 13 weeks.

**Conclusion:** These data demonstrate that perturbed hemodynamic factors triggered by lung microvascular arteriolar loss play a requisite role in perpetuating endothelial injury in experimental PAH, leading to persistent arterial EC injury and disease progression. Importantly, vascular loss, arterial remodeling and PH are reversible once the cycle of perturbed hemodynamics and EC injury is broken by unilateral lung banding.

## 2. Introduction

Pulmonary arterial hypertension (PAH) is a disease of the lung arterial bed, resulting in the progressive loss of effective microvascular area leading to increases in pulmonary vascular resistance (PVR) and pulmonary hypertension (PH).^1, 2^ However, the mechanisms underlying the development of PAH are hotly debated.^1^ Recently, there has been considerable interest in the proliferative changes that are responsible for complex arterial remodeling which represents a hallmark feature of this disease.^3-6^ Mounting evidence supports dysregulated vascular cell growth, resulting in narrowing and occlusion of distal lung arteries, as a dominant mechanism in the pathogenesis of PAH.^3-6^ This ‘cancer-like’ hypothesis represents an attractive paradigm that leverages a wealth of mechanistic insights into pathways of growth dysregulation from cancer biology.^7-9^ Many parallels between cancer and hyperproliferative and apoptosis-resistant cells in proliferative arterial lesions in PAH have been identified, including upregulation of cell growth and survival pathways^10, 11^ and epigenetic and DNA repair mechanisms, which have led to a number of novel therapeutic strategies that are being explored in preclinical models and early clinical trials (Imatinib, BDR4 inhibitor, etc.).^6, 12, 13^

However, unlike most cancers, dysregulated vascular cell growth in PAH is triggered in response to external signals, in particular ongoing EC injury likely caused by increased intimal shear stress. Indeed, Abe et al. demonstrated that occlusive arterial remodeling could be abrogated by reducing pulmonary blood flow in the SU5416-chronic hypoxia (SUHx) rat model of severe PAH.^14^ Moreover, unilateral pulmonary artery banding could reverse established complex arterial lesions when banding was delayed to 5 weeks post. These findings suggest that dysregulated vascular cell growth is dependent on severe hemodynamic perturbations, due to increased luminal shear stress, yet it is unclear what causes the hemodynamic abnormalities in the first place in this model. Abe speculated that this may be due to hypoxia-induced vasoconstriction; however, the development of a severe PAH phenotype in response to SU alone, even in absence of hypoxia in immunodeficient nude rats^15^, and in a hyper-responsive substrain of Sprague-Dawely (SD) rats^16^, points to the involvement of other mechanisms.

Lung endothelial apoptosis is well recognized as a central trigger for the initiation of PAH in experimental models and human disease.^17^ It has recently been suggested EC apoptosis could result directly in the degeneration and loss of fragile precapillary arterioles^1, 18^ leading to widespread microvascular pruning as a mechanism for the initial increases in pulmonary arterial pressure (PAP) and PVR in this model. Moreover, since the lung must accommodate the entire cardiac output, any reduction in lung arteriolar area would necessarily produce microcirculatory hemodynamics changes, in particular an increase in shear forces in the remaining lung microvasculature. Increased intimal shear stress is well known to cause ongoing endothelial injury in PAH associated with congenital heart disease and left to right shunting^19^. Therefore, in this study we assessed the relationship between lung arteriolar pruning, pulmonary blood flow and the development of PH in the SUHx rat model of PAH. Our findings point to loss of lung arteriolar microcirculation as a major mechanism in the onset of PH after SU-induced endothelial injury. Moreover, we show that the resultant alterations in shear forces within the lung microcirculation play a requisite role in perpetuating endothelial injury and arteriolar loss, resulting in persistent arterial remodeling and PH. Importantly, arterial pruning and PH were reversible once the vicious cycle of perturbed lung hemodynamics and ongoing EC injury was interrupted by unilateral lung banding.

## 3. Methods

Detailed methods are available in the online supplement. All animal care and study protocols were approved by the University of Ottawa Animal Care Committee and conducted according to the guidelines from the Canadian Council for Animal Care. All procedures were according to the guidelines from the NIH Guide for the Care and Use of Laboratory Animals.

### 3.1 Rat SU5416 chronic hypoxia (SUHx) model of PAH

Male Sprague Dawley (SD, Harlan laboratories, IN, USA) rats weighing 150-200 g were used for this study. PAH was induced by a single subcutaneous injection of SU5416 (SU:3-(3,5-dimethyl-1H-pyrrol-2-ylmethylene)-1,3-dihydroindol-2-one) (Tocris, Bristol, United Kingdom) in 0.5% carboxymethyl cellulose followed by 3-weeks chronic hypoxia (9-10% O_2_), as previously described.^16, 20^ Following completion of hypoxia treatment, the rats were housed under normoxic.

### 3.2 Left main pulmonary artery banding (LPAB)

SUHx rats were subjected to LPAB or sham procedure at 1-week prior (prevention) or 5 weeks after (reversal) SU injection as described earlier and supplemental material.^14^

### 3.3 Non-invasive assessment by echocardiography

Echocardiography was performed at the end study using the Vevo2100 ultrasonography system (VisualSonics, ON, Canada) as described earlier.^20^

### 3.4 Measurement of RVSP and RV hypertrophy

RVSP was measured using high-fidelity pressure catheters (Transonic-Scisense Inc., ON, Canada) as described earlier.^20^

### 3.5 Lung perfusion and microCT analysis of pulmonary circulation

microCT analysis of lungs were performed to assess pulmonary circulation as described earlier.^21^ Samples were imaged using a desktop micro-CT (SkyScan 1272, Bruker microCT, Kontich, Belgium) and analyzed as described previously.^21^

### 3.6 Lung histological measurements

Paraffin sections were then utilized for histology and immunostaining as described previously^20^ and in supplemental material. Percent Medial wall thickness = ((distance between the internal and external lamina × 2)/external diameter) × 100. For total vessel count, all the vessels (<100μm) were counted from the 10 random fields and divided in grade-1 (30-80% occlusion) and grade-2 (>80% occlusion).

### 3.7 Immunostaining of lung sections

Immunohistochemistry for cleaved caspase-3 Cell Signalling Technologies, Cat# 9661S) and Von Willebrand Factor (vWF, Abcam, Cat# ab6994) was performed as described earlier.^20, 22^ Immunofluorescence staining for Ki67 (Abcam, Cat# ab15580) and CD144 (R&D systems, Cat# BAF1002) was performed as described in supplemental material.

### 3.8 Statistical analysis

Data are represented as mean±SEM unless otherwise stated. Statistical analysis was performed using GraphPad Prism 9.0 (GraphPad Software, Inc. CA, USA). For statistical comparisons, Student’s t-test or One-Way ANOVA (>2 groups) were performed followed by Dunnett or Sidak multiple comparison test with significance level of p<0.05. Paired t-test was used for comparison of repeated measures.

## 4. Results

### 4.1 Development of PH is associated with persistent pulmonary arteriolar EC apoptosis and progressive arterial pruning

We made serial assessments of RVSP, RV hypertrophy, EC apoptosis, pulmonary vascular density and pulmonary arterial remodeling at 1, 3, 7, 21 and 49-days post-SU injection in the SUHx rat model of PAH. There was a progressive increase in RVSP and Fulton index (RV hypertrophy) with the significant changes as early as day-7 (Fig. 1A and 1B). This was associated with increased cleaved caspase-3, localized by immunostaining to arterioles, which was significantly increased at day-3 and remained elevated throughout the 49 days of follow up (Fig. 1C and 1D). Furthermore, ongoing EC apoptosis was mirrored by a reduction in the number of arterioles (<50 μm in diameter) per unit area in lung sections beginning day-3 post-SU (Fig. 1E and Supplemental Fig. 1A). Importantly, the reduction in vessel counts strongly correlated with the proportion of cleaved caspase-3 positive intimal cells in pulmonary vessels (Supplemental Fig. 1B). The increase in medial wall thickness (Fig 1F and Supplemental Fig. 2) and evidence of arteriolar occlusive remodeling (Fig. 1G) was not seen until 7-days and 49-days post-SU injection, respectively.

**Figure 1:**
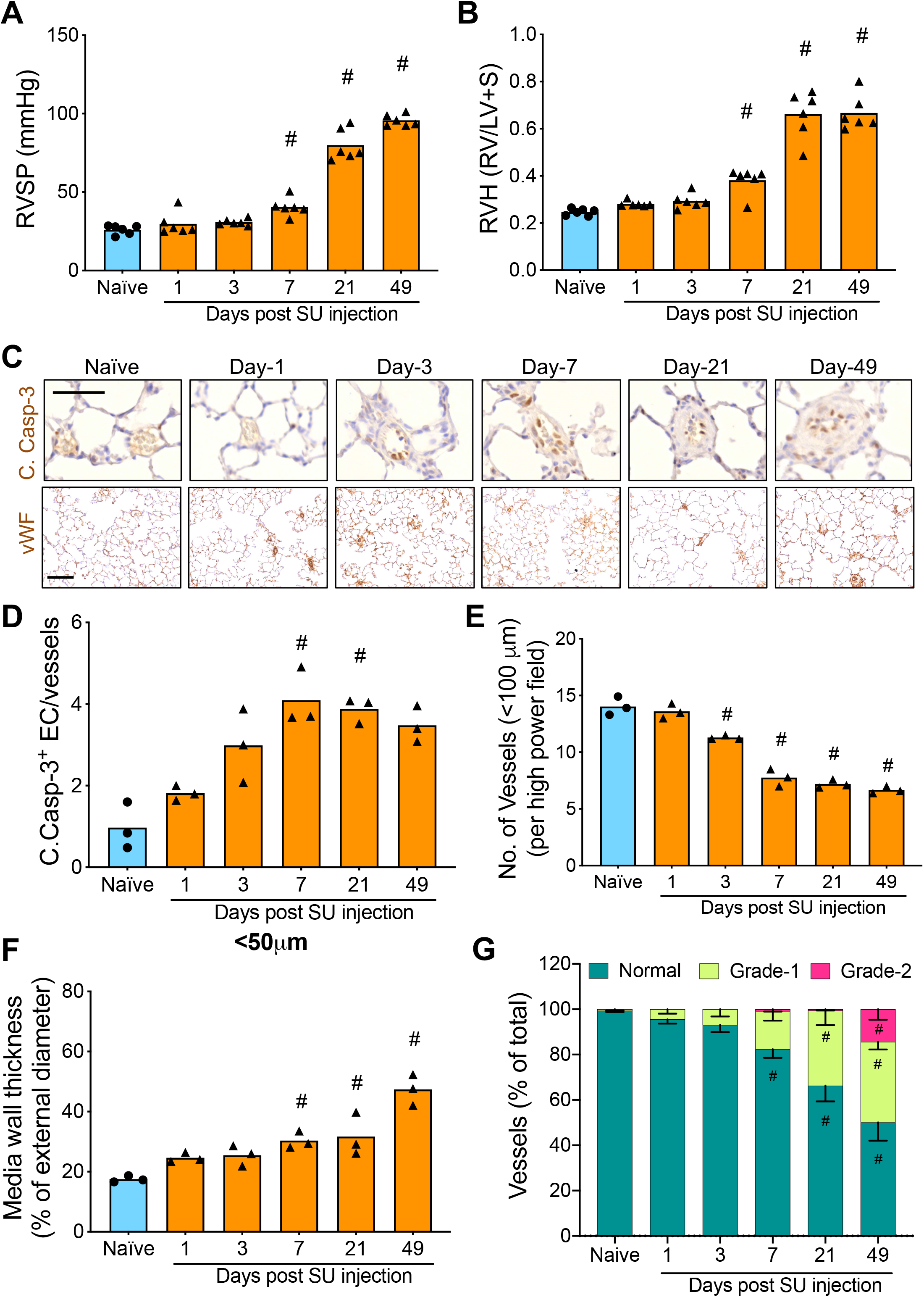
Progressive PAH in SUHx model. Bar graphs demonstrating **A)** RVSP and **B)** RV hypertrophy of control and SUHx treated male SD rats at different time-points post-SU. N=6 per group, bar graph showing mean, ^#^p<0.05 vs naïve rats. **C)** Representative micrograph images demonstrating cleaved caspase-3 staining in pulmonary vasculature and von-Willebrand Factor (vWF) staining in left lung sections. Scale bar = 50 μm. **D)** Bar graph showing quantification of cleaved caspase-3 positive intimal cells in small pulmonary arteries at different time-points post-SU in rat SUHx model. N=3 per group, bar graph showing mean, ^#^p<0.05 vs naïve rats. **E)** Number of small arteries (<100 μm diameter) in lungs of SUHx subjected male SD rats at different time-points post-SU, N=3 per group, bar graph showing mean±SEM, ^#^p<0.05 vs naïve rats. **F)** Media wall thickness of <50 μm diameter vessels of male SD rats subjected to SUHx model at different time-points post-SU, N=3 per group, bar graph showing mean, ^#^p<0.05 vs naïve rats. **G)** Vascular occlusion in lungs of SUHx subjected male SD rats at different time-points post-SU, N=3 per group, bar graph showing mean±SEM, ^#^p<0.05 vs naïve rats.

To further assess the impact of EC apoptosis on pulmonary circulation, we performed micro-CT analysis of the functional arterial bed at 4- and 7-weeks post injection of SU or CMC-vehicle followed by a 3-week exposure to hypoxia (Fig 2A). Four weeks was chosen as the earliest time point to avoid the potentially confounding effects of hypoxia. A significant reduction in arterial volume was observed in hypoxia together with CMC-vehicle treated rats at 4 weeks timepoint compared to naïve rats (Fig 2B-C), associated with a modest increase in RVSP (Fig 2D). However, both arterial volume and PH recovered at the 7 weeks timepoint in hypoxia alone rats (Fig 2C and D). In contrast, there was a marked reduction in arterial volume in SUHx group at 4 weeks that persisted until 7-week timepoint (Fig 2A-C). Of note, arteriolar pruning was limited to smaller arteries of <200μm diameter with minimal reduction in arterial volume in larger arteries (Fig 2B). Similarly, RVSP in SUHx group was markedly elevated at 4 weeks post-SU and remained elevated at 7-week post-SU (Fig 2D). RVSP and arterial volume were not significantly affected at in CMC-treated normoxic rats at 4- or 7-weeks post-SU. Next, we examined the correlation between loss of vessel volume and RVSP. Overall, the relationship was highly nonlinear with no significant increase in RVSP until there was an ∼75% loss in arterial volume (Fig. 2E). However, after this threshold was achieved, a strong correlation between loss of arterial volume and increase in RVSP was observed. This is consistent with the ability of the lung to accommodate the cardiac output at relatively normal pressures until about 80% of the arterial bed has been compromised, at which point thus produces hemodynamic perturbations in the remaining lung vasculature which are similar to those induced by a large left to right arterial shunt.

**Figure 2:**
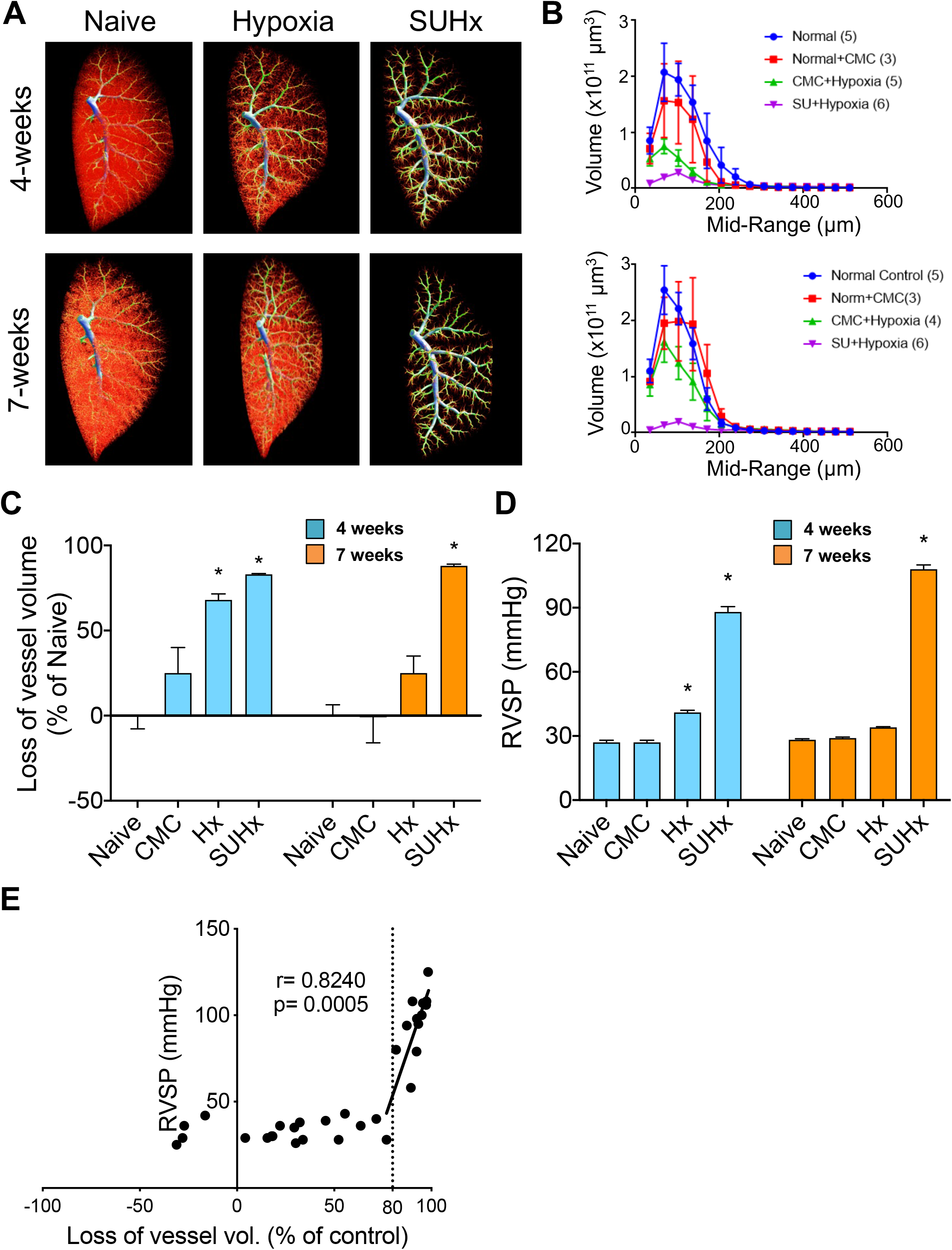
Increased hemodynamic load in SUHx model is associated with loss of pulmonary vascular volume. **A)** Representative microCT images and **B)** line graph demonstrating pulmonary vascular volume in naïve, CMC-vehicle, hypoxia or SUHx subjected male SD rats at 4- and 7-week time-points post-SU. N=3-6 per group, values represent mean±SEM. **C)** Bar graph showing estimated loss of volume (calculated as a percentage of naïve) in response to CMC, hypoxia or SUHx at 4- and 7- week time-points post-SU. N=3-6 per group, values represent mean±SEM. *p<0.05 vs naïve rats. **D)** Bar graph showing RVSP in naïve, CMC, hypoxia or SUHx at 4- and 7-week time-points post-SU. N=3-6 per group, values represent mean±SEM. **E)** Dot-plot demonstrating correlation between loss of vascular volume and RVSP. Line represents correlation between loss of vascular volume and RVSP for samples with >80% loss of vascular volume. r shows Pearson coefficient.

### 4.2 Hemodynamic offloading by LPAB prevented vascular EC apoptosis and abolished the loss of vascular volume in left lung

Next, we sought to determine the effect of restricting lung blood flow on persistent endothelial injury and arteriolar loss in this model. Unilateral hemodynamic offloading was achieved by left pulmonary artery banding (LPAB) as described previously.^14^ LPAB or sham surgery (Sham) was performed 1- week prior to SU and RVSP, arterial volume (microCT) and EC apoptosis was assessed at 8 weeks post SU (Fig. 3A). There were no significant differences in RVSP and RVH between the sham and LPAB groups (Fig. 3B-C). LPAB prevented ongoing endothelial injury in the banded lung, whereas persistent cleaved caspase-3 immunostaining was evident in arteriolar ECs in the non-banded right lung (Fig. 3D). Similarly, vascular cell proliferation, evidenced by Ki67 immunostaining, was seen in intima and media of small arterioles in the right lung, but was absent in the banded left lung (Fig. 3E, supplemental Fig 3A). Consequently, a significant increase in partially and completely occluded arterioles was observed in the nonprotected right lungs of LPAB rats, whereas the left lungs were largely free of occlusive arterial remodeling (Fig. 3F). Furthermore, microCT analysis showed preservation of arterial volume in the protected left lungs of LPAB rats (Fig. 3G-H). Notably, the normal left lung arterial volume is only 40% of total lung arterial volume (Supplemental Figure 3B-C). However, after LPAB, the left lung arterial volume was similar to that of the right lung, whereas in sham-operated rats the arterial volume in the left lung was markedly reduced (Fig 3G-H). Overall, these studies demonstrated that hemodynamic offloading prevented protracted endothelial injury, occlusive arterial remodeling and loss of arterial volume in the banded left lung of rats in SUHx model.

**Figure 3:**
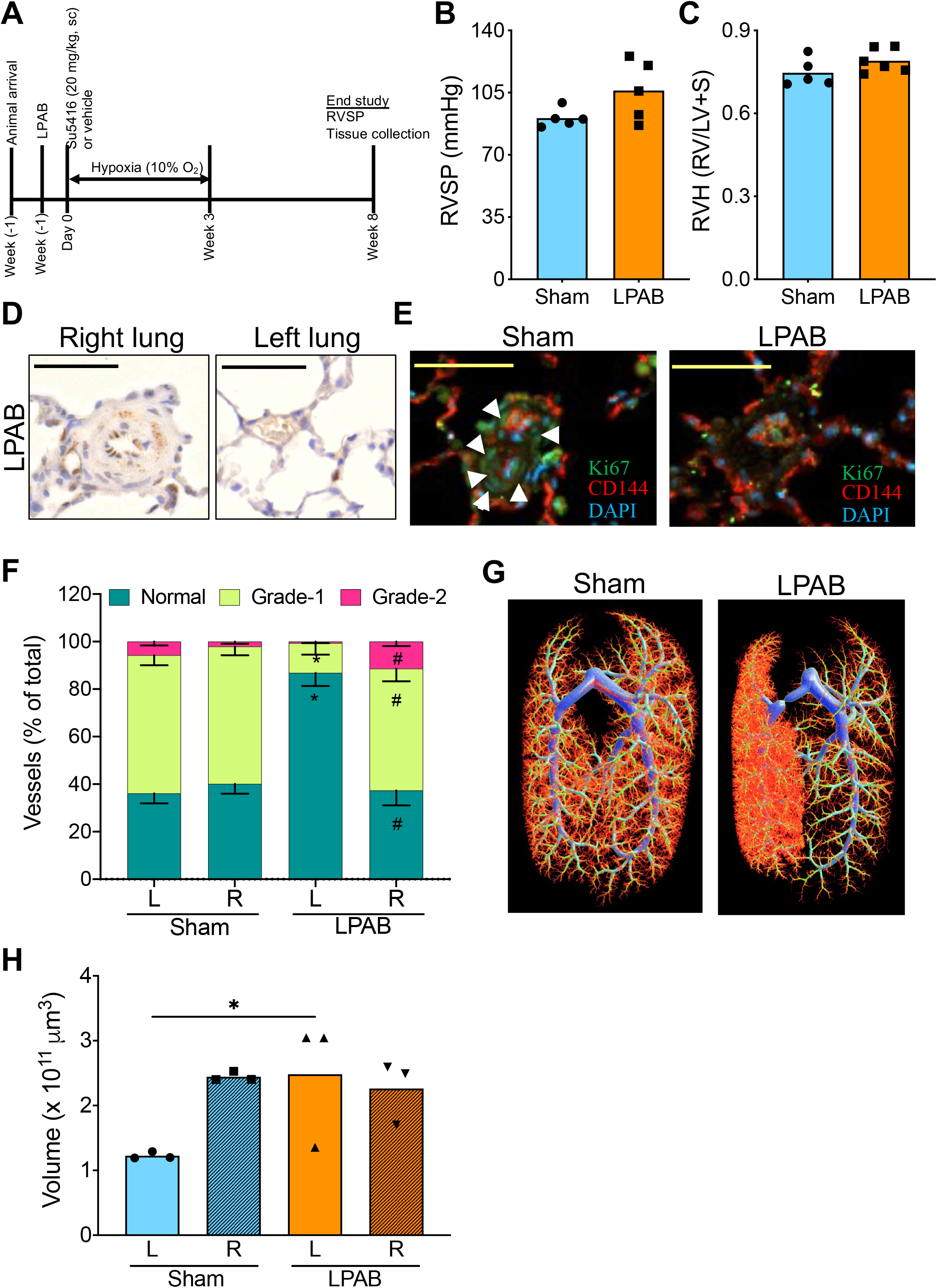
Hemodynamic offloading by LPA banding prevents pulmonary vascular remodeling in SUHx model. **A)** Schematic diagram of experimental timeline to study the effect of prevention (at 2 weeks before SU) LPA banding in SUHx model. **B)** RVSP and **C)** RV hypertrophy of sham operated or LPAB male SD rats subjected to SUHx. N=5-6 per group, bar graph showing mean. **D)** Representative micrograph images demonstrating cleaved caspase-3 staining in pulmonary vasculature of left and right lung of SUHx treated LPAB rats at 8 weeks post-SU. Scale bar = 50 μm. **E)** Representative micrograph images demonstrating immunofluorescence staining for Ki67 (proliferation marker) and CD144 (EC marker) in pulmonary vasculature of left and right lung of SUHx treated rats following LPAB. Scale bar = 50 μm. **F)** Vascular occlusion in left and right lung of LPAB or sham operated male SD rats subjected to SUHx at 8-weeks post-SU, N=6 per group, bar graph showing mean±SEM, *p<0.05 vs left lung of sham group, ^#^p<0.05 vs left lung of LPAB group. **G)** Representative microCT images and **H)** bar graph showing vascular volume of left and right lung of SUHx treated LPAB rats at 8 weeks post-SU. N=3 per group, bar graph showing mean±SEM, *p<0.05.

### 4.3 Unilateral flow restriction at 5 weeks post-SU alleviates ongoing arterial endothelial injury and reverses arterial pruning in the protected lung

We then investigated whether delayed hemodynamic offloading at 5 weeks post-SU (reversal LPAB) could alleviate ongoing EC injury and restore arterial volume after widespread arterial pruning has already occured in this severe PAH model. We randomized the rats to delayed LPAB or sham surgery and then assessed EC apoptosis, arterial remodeling and volume (microCT) at 8- or 10-weeks post-SU (i.e., 3 or 5 weeks post banding; Fig 4A). Again, we did not observe any significant differences in RVSP and RVH between LPAB and sham groups at 8 weeks post SU (Fig 4B-C). Interestingly, the number of cleaved caspase-3 positive ECs was markedly lower in left lung of LPAB rats, while robust immunostaining was observed the right lung of the LPAB group and in both lungs of sham rats (Fig 4D and Supplemental Fig. 4). While only partial improvement in arterial volume was evident at 8-weeks post-SU (3 weeks post-LPAB), there was a marked improvement in arterial volume in left lung at 10 weeks post-SU (5 weeks post-LPAB) (Fig 4F-G). These data show that hemodynamic offloading by LPAB alleviates endothelial injury in established PAH and allows for progressive microvascular repair and restoration of the lung arterial bed.

**Figure 4:**
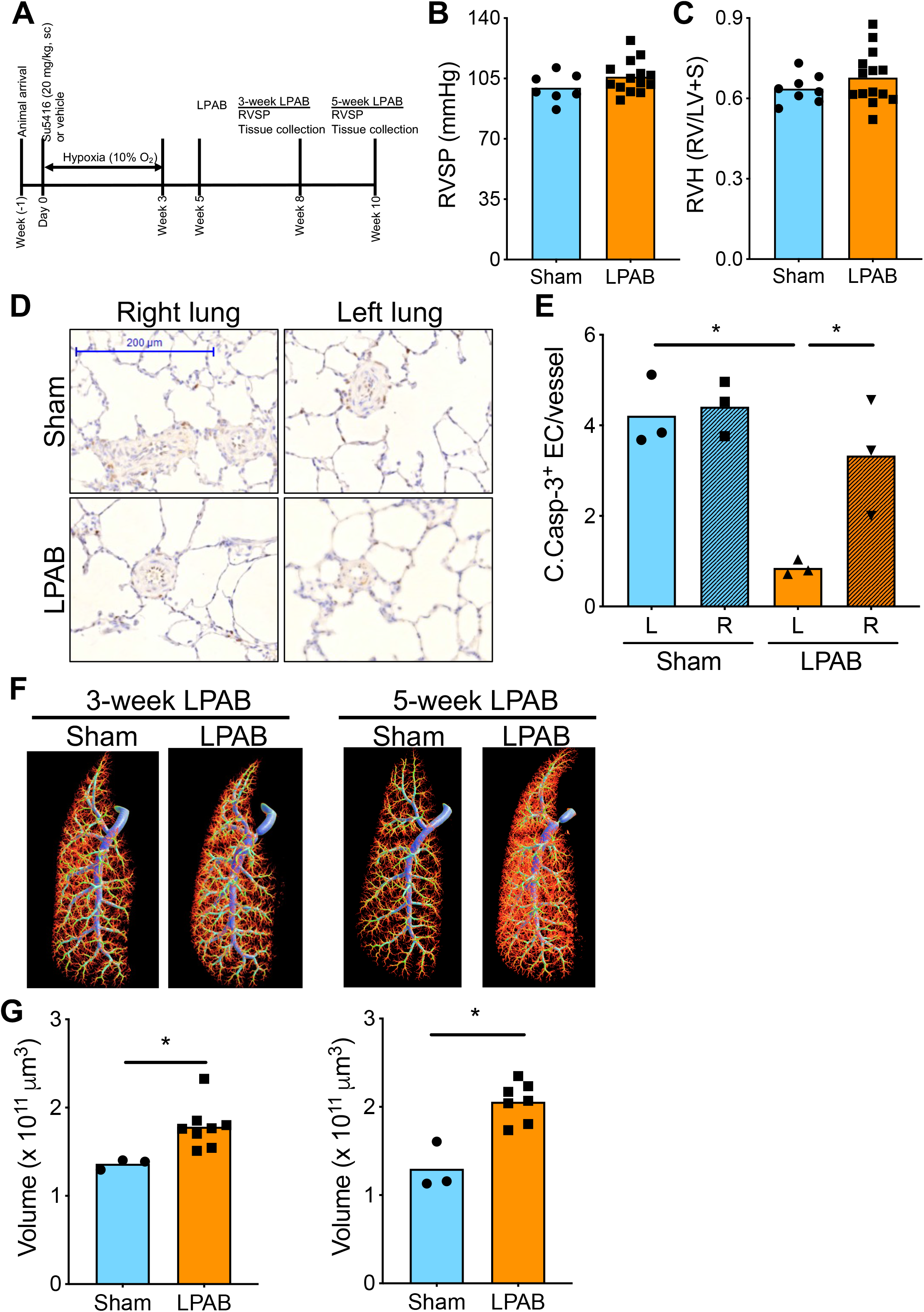
Hemodynamic offloading by LPA banding reverses pulmonary vascular remodeling in SUHx model. **A)** Schematic diagram of experimental timeline for the study of the effect of reversal (at 5 weeks post-SU) LPA banding in SUHx model. **B)** RVSP and **C)** RV hypertrophy of sham operated or LPAB male SD rats subjected to SUHx. N=4-6 per group, bar graph showing mean. **D)** Representative micrograph images and **E)** bar graph demonstrating cleaved caspase-3 staining in pulmonary vasculature of left and right lung of SUHx treated LPAB rats at 8 weeks post-SU (3 weeks LPAB). N=3 per group, bar graph showing mean±SEM, *p<0.05. **F)** Representative microCT images demonstrating pulmonary vasculature and **H)** bar graph showing quantification of vascular volume of left lung of SUHx treated LPAB rats at 8- or 10-weeks post-SU (3 or 5 weeks LPAB, respectively). N=3-8 per group, bar graph showing mean, *p<0.05.

### 4.4 Partial improvement of PH after late removal of LPAB

Finally, we assessed whether unbanding of the left pulmonary artery would have a beneficial effect on pulmonary hemodynamic in the SUHx PAH model. SUHx or sham rats were randomized to receive sham or LPAB surgery at 5 weeks post-SU, as above. Five weeks after surgery (10 weeks post-SU), the LPAB rats underwent an unbanding or sham procedure, and RVSP, RVH and RV function were assessed 3 weeks later (i.e., 13 weeks post SU). As shown in Fig 5A, 3 groups were studied: 1), sham surgery at 5 weeks and sham surgery at 10 weeks (Sham-Sham); 2) LPAB surgery at 5 weeks and sham surgery at 10 weeks (LPAB-Sham) and 3): LPAB surgery at 5 weeks and LPAB removal surgery at 10 weeks (LPAB-Unband). Once more, there were no significant differences in RVSP immediately prior to unbanding surgery (10-weeks) between the three groups, (Fig 5B). However, 3 weeks after unbanding of the left pulmonary artery, there was a significant reduction in RVSP in the LAPB-unband group compared to either the LPAB-Sham or the Sham-Sham groups (Fig 5C). Since a hemodynamic assessment was made at both 10 and 13 weeks for each animal, a paired analysis was performed (Fig D-F) which again showed an ∼36.8% reduction in RVSP in the unbanded animals. Interestingly, there was also a significant albeit modest improvement (∼16.7%) in the Sham-Sham group, consistent with a previous report showing some spontaneous improvement in pulmonary hemodynamics at later timepoints in the SUHx model.^16^ Nonetheless, the LPAB-Sham group, which is the more appropriate control, showed no spontaneous hemodynamic improvement, and even a tendency in most animals toward progression, possibly reflecting more severe disease in the unprotected right lung due to the relative increase in blood flow. Moreover, a significant reduction in Fulton index was observed in LPAB-unbanded group compared to LPAB-sham group (Fig 5G). This improvement was associated with an increase in cardiac output (CO) and reduction in PVR in LPAB-unbanded group compared to both sham-sham and LPAB-sham groups (Fig 6A and B). There was also an improvement in right ventricular function as assessed by stroke volume and fractional shortening in LPAB-unbanded group compared to LPAB-sham group (Fig 6C and 6D). No significant difference in heart rate was observed between any of the three groups (Fig 6E). Overall, these data show that reintroduction of the protected left lung vascular bed into the circulation after hemodynamic offloading was sufficient amount to result in a meaningful improvement established PAH and RV function.

**Figure 5:**
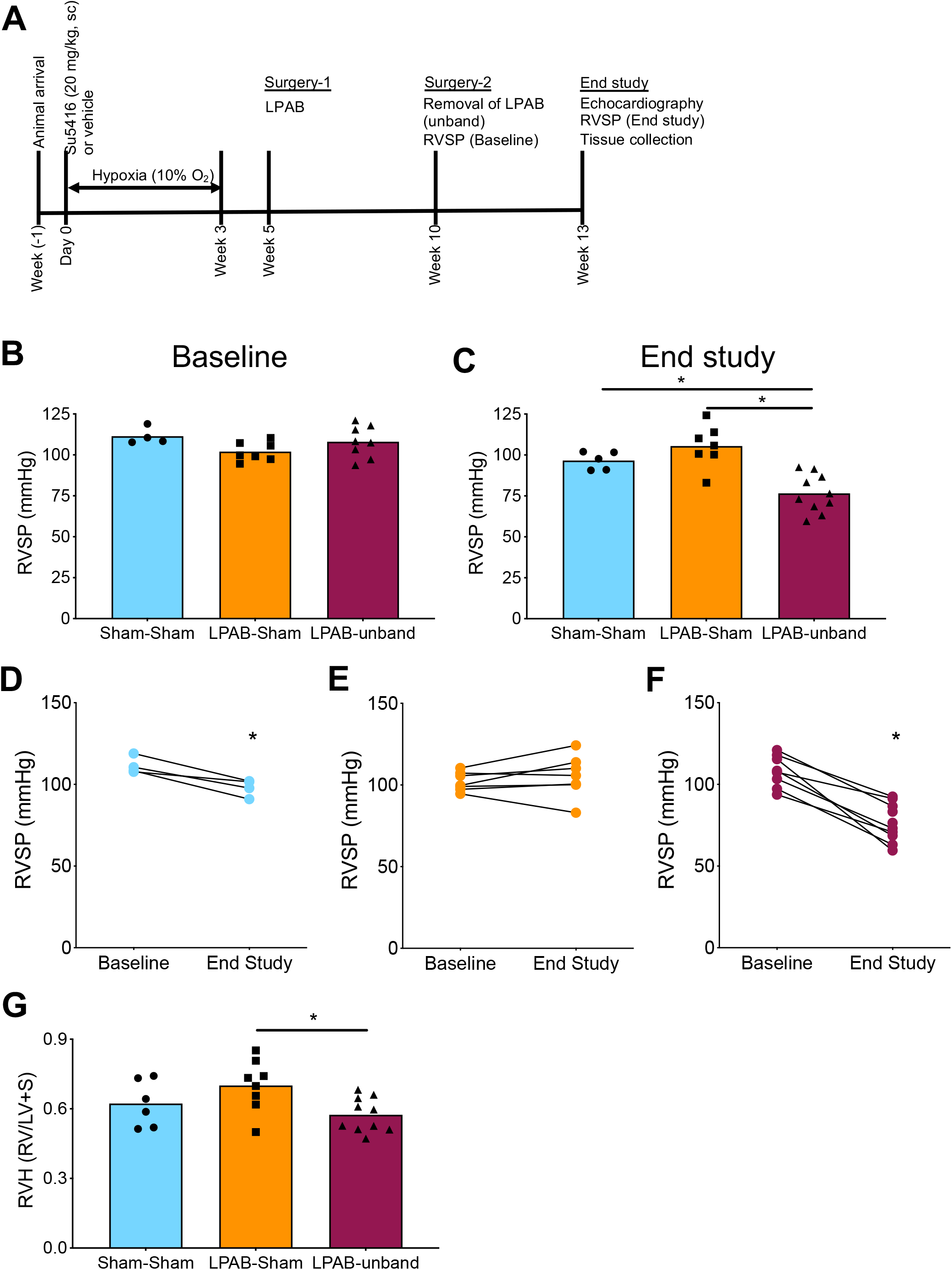
Hemodynamic offloading by LPA banding reverses PAH in SUHx model. **A)** Schematic diagram of experimental timeline to study the effect of reversal LPA banding on PAH phenotype in SUHx model. **B)** Baseline (10 weeks post-SU) and **C)** end study (13 weeks post-SU) RVSP of rats subjected to SUHx followed by sham surgery at 5-weeks and 10-weeks post-SU (sham-sham group), LPAB surgery at 5-weeks and sham surgery at 10-weeks post-SU (LPAB-sham group) or LPAB surgery at 5-weeks and unband surgery at 10-weeks post-SU (LPAB-unband group). N=4-10 per group, bar graph showing mean, *p<0.05. Change in RVSP of individual rats from baseline (10 weeks post-SU) to end study (13 weeks post-SU) in **D)** sham-sham group, **E)** LPAB-sham group and **F)** LPAB-unband group. *p<0.05. **G)** Bar graph showing RV hypertrophy of sham-sham, LPAB-sham and LPAB-unband groups of rats. N=6-10 per group, bar graph showing mean, *p<0.05.

**Figure 6:**
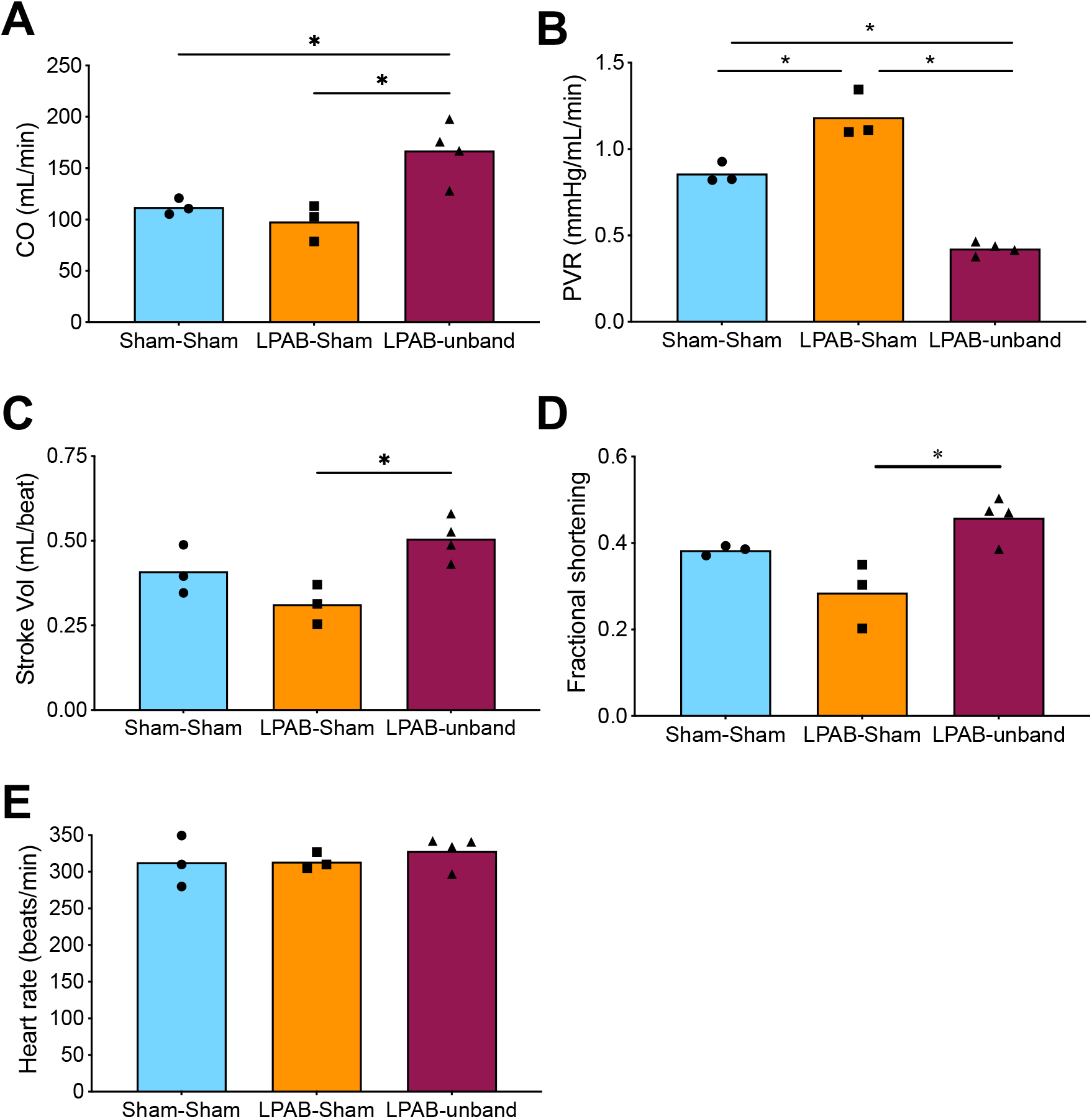
LPA banding mediated improvement in pulmonary vasculature improves cardiopulmonary function in the SUHx model. Bar graphs showing **A)** Cardiac output (CO); **B)** pulmonary vascular resistance (RVSP/CO); **C)** stroke volume, **D)** fractional shortening [(RVIDd-RVIDs)/RVIDd] and **E)** heart rate of sham-sham, LPAB-sham and LPAB-unband groups of rats. N=3-4 per group, bar graph showing mean, *p<0.05.

## 5. Discussion

PAH is an incurable disease that afflicts people of all ages, including children and young adults, mainly affecting women^23^, and is usually progressive despite all available therapies, ultimately leading to right heart failure and death or lung transplantation.^24^ While EC apoptosis is widely regarded as a trigger for the development of severe PAH, it is unclear how this leads to and complex vascular remodeling and protracted PH over the long term. We mapped the time course of EC apoptosis in the SUHx severe PAH model and established the role arterial pruning and perturbed pulmonary blood flow in both the onset and persistence of disease. For the first time we demonstrate that pulmonary EC apoptosis persisted for at least 8 weeks, far beyond the period that could reasonably be attributed to the direct pharmacologic effects of a single dose of SU, even recognizing that it is delivered in a depot formulation with CMC and that inhibitory intracellular concentrations of SU may persist for about 72 hours beyond its plasma half-life.^25^

We also showed that persistent EC apoptosis was dependent on pulmonary blood flow, consistent with ongoing endothelial injury induced by perturbed hemodynamics in this model. Therefore, we quantified changes in effective arterial volume during the onset of PH in this model and related this to the time course of EC apoptosis and proliferation. Using a highly quantitative and optimized micro-CT approach that had been validated extensively,^21^ we determined arterial volumes in the weeks after induction of EC injury with SU. We found surprisingly rapid onset of pruning lung arterioles, mainly <200 microns, which reached near maximal at 4 weeks, and closely mirrored the onset of PH. Although there was evidence of vascular cell proliferation during this period, as well as modest increased medial thickness in small and medium-sized arterioles, there was no evidence of occlusive arterial remodeling during the first 3 weeks, which is consistent with other reports showing that complex lesions are only seen after 5 weeks.^26^ In contrast, a significant decrease in lung vascular density was apparent as early as 1 week post SU, raising the possibility that the direct effects of EC apoptosis, specifically the degeneration and pruning of the fragile distal arteriolar bed, was the cause of the initial increases in pulmonary arterial pressure and resistance in this model.

Since the lung must accommodate the entire cardiac output, marked reductions in lung microvascular cross-sectional area necessarily result in increased luminal shear stress at the intimal surface. This is analogous to increases in shear stress associated with left to right shunts in the context of congenital heart disease.^1, 19^ In both cases there is a mismatch between lung blood flow and arterial cross-sectional area resulting high shear forces which induce lung endothelial injury.^1^ In patients with congenital heart disease, this mechanism is thought to underly the development of PAH and ultimately Eisenmenger’s syndrome. Indeed, complex arterial remodeling was initially described in lung specimens from Eisenmenger patients and these lesions are identical to those seen in other forms of PAH.^27^ Interestingly, with the exception of the SUHx model, the only other preclinical models that reliably reproduced complex arteriopathy required increasing lung blood flow by creating a left to right shunt^28^ or by unilateral pneumonectomy^29^ together with endothelial injury. This underscores the importance of perturbations in hemodynamic factors, in particular lung blood flow and increase luminal shear forces, in the structural arterial changes of PAH.

The mechanisms by which high lung microvascular shear stress contributes to the development of PAH are still unclear. In CHD, elevated pulmonary blood flow by itself rarely results in PAH^30^, and it thought that other factors, such as endothelial dysfunction or elevated pulmonary arterial pressure, are required.^31^ Indeed, both of these factors are present in the SUHx model which is initiated by inhibition of VEGF signaling resulting in EC apoptosis and is characterized by severe PH.^17^ Since LPAB will not only reduce pulmonary blood flow, but also arterial pressure as a result of pressure loss across the flow limiting stenosis, it is not possible to separate their relative contributions to the protection of the lung arterial bed. Nonetheless, there is an abundant evidence to support the role of pathological levels of intimal shear stress in the pathogenesis of CHD related PAH, including promoting endothelial dysfunction^32^ and abnormalities in EC adaptation.^33, 34^ However, the role of pathological intimal shear stress in inducing EC injury and apoptosis is less clear. Indeed, high intimal shear stress has been shown to be protective against EC apoptosis and atherosclerosis in large systemic arteries, and disturbed flow has been implicated in EC injury.^35, 36^ In the pulmonary circulation, a relative increase in blood flow will result in a disproportionate increase in shear forces in the distal lung arteriolar bed^37, 38^, which is plays such a central role in PAH, as well as disturbed flow patterns at branch points^39, 40^, both of which could produce EC injury. Indeed, we showed sustained increases in EC cleaved caspase 3 expression over the course of PAH progression in the SUHx model which was dependent of perturbed pulmonary hemodynamics and was normalized after LPAB. To our knowledge, this is the first demonstration of ongoing EC apoptosis attributable to hemodynamic shear forces in PAH, and points to this as a key mechanism driving persistent arteriolar pruning and complex remodeling in this disease.

This essential role of hemodynamic perturbation and increase shear stress in perpetuating EC injury and vascular loss was confirmed by unilateral pulmonary artery banding in the SUHx model. Remarkably, not only did this prevent and reverse complex arterial remodeling, as had been previously reported^14^, but it also mitigated the effects of SU and hypoxia on EC apoptosis and arteriolar loss. This was further exemplified by studies in which the pulmonary artery banding was relieved. Indeed, when the left lung was banded after PAH had been fully established, removal of the flow restriction resulted in a significant improvement in overall pulmonary hemodynamics. This indicates that the improvements in lung arteriolar density and structure that were achieved by LPAB are functionally meaningful, and when the protected lung arterial bed was reintegrated into the pulmonary circulation, there was a sustained reduction in pulmonary arterial pressure and resistance. However, the hemodynamic improvement was only partial, likely because the left lung represents only <40% of the total lung arterial volume, so that even if there was a full recovery in vascularity, this might not be sufficient to fully accommodate the normal cardiac output. Moreover, given the relatively lower vascular resistance in the protected left lung, it is likely that after debanding, there would be a disproportionate redistribution of blood flow to the left lung resulting in a return of pathological increases in shear forces, leading to recurrence of arterial pruning and remodeling over time.

Our study has some notable limitations. First, the earliest timepoint that arterial pruning was assessed was 4 weeks after administration of SU since we wanted to avoid the confounding effects of exposure to hypoxia. However, in a subsequent study, significant pruning of the arterial bed was apparent as early as 1 week and was near maximal at 3 weeks, closely mirroring the increases in RVSP at these time points.^41^ Second, the relative blood flow was not measured between the left and right lungs before and after banding. Nonetheless, in our pilot experiments we found that a >70% narrowing was necessary to produce vascular protection after LPAB, which is consistent with the need for flow restricting stenosis. Also, we were unable to confirm the degree to which the stenosis was relieved after the band was removed. Due to local reaction and fibrosis, the reopening of the left pulmonary artery was likely gradual and may have variable form on animal to another. Nonetheless, we were able to show significant and important overall hemodynamic improvements by 3 weeks after unbanding.

Therefore, these data show that lung arterial pruning and structural changes were fully reversible when the cycle of hemodynamic perturbation and endothelial injury was interrupted by pulmonary artery banding. These findings have some very provocative implications. First, they support the idea that EC apoptosis plays a direct role in the initiation of PAH by causing degeneration and drop-out of distal lung arterioles. Second, they suggest that arterial proliferative lesions occur in response to ongoing endothelial injury and apoptosis, driven by hemodynamic perturbation, in particular increases in shear forces, as a result of critical reduction in the effective area of the lung microcirculation. Third, they show that ongoing EC apoptosis and its consequences, specifically lung microvascular loss and complex arterial lesion formation, are preventable and reversible by restricting blood flow to the lung microcirculation. While this does not rule an important role for complex arterial remodeling in the progression of established PAH, it clearly demonstrates that this can be obviated by interrupting the underlying feed-forward cycle of endothelial injury and hemodynamic abnormalities. These new insights suggest novel therapeutic directions aimed at mitigating endothelial injury and promoting vascular repair as potentially effective treatment strategies for this devastating disease.

### 6. Funding

This research was supported by a grant from Actelion Pharmaceuticals US, Inc. ENTELLIGENCE Young Investigator Program to KRC. This study was funded by a Catalyst grant from Canadian Institute of Health Research to DJS and KRC (Funding Reference# SVB-145555). KRC received Scholar award from Canadian Vascular Network and Heart and Stroke Foundation of Canada.

## 7. Acknowledgements

We would like to thank Xiaoxue Wen and Katelynn Rowe for their technical support in animal procedures.

