## Supplemental data for "Vicious cycle of hemodynamic perturbation and endothelial injury in development and progression of pulmonary arterial hypertension"

### Short Title: Arteriolar loss and hemodynamic stress in PAH progression

### Yupu Deng^1^*; Ketul R. Chaudhary^1,2,3^*; Anli Yang^1^; Kirishani Kesavan^3^; Liyuan Wang^1,2^, Kevin Chathely^1^, Duncan J. Stewart^1,2^

### ^1^Sinclair Centre for Regenerative Medicine, Ottawa Hospital Research Institute, Ottawa, Canada

### ^2^Department of Cellular and Molecular Medicine, Faculty of Medicine, University of Ottawa, Ottawa, Ontario, Canada

^3^Department of Physiology and Biophysics, Faculty of Medicine, Dalhousie University, Halifax, Nova Scotia, Canada

*** Both authors contributed equally to the work.**

**Address for Correspondence:**

Duncan J. Stewart, M.D.

CEO and Scientific Director, Ottawa Hospital Research Institute

501 Smyth Road, Ottawa, Ontario, K1H 8L6, Canada

**Category**: Original research article

**Methods:**

All animal care and study protocols were approved by the University of Ottawa Animal Care Committee and conducted according to the guidelines from the Canadian Council for Animal Care. All procedures were according to the guidelines from the NIH Guide for the Care and Use of Laboratory Animals.

**Rat SU5416 chronic hypoxia (SUHx) model of PAH**

Male Sprague Dawley (SD, Harlan laboratories, IN, USA) rats weighing 150-200 g were used for this study. PAH was induced by a single subcutaneous injection of SU5416 (SU:3-(3,5-dimethyl-1H-pyrrol-2-ylmethylene)-1,3-dihydroindol-2-one) (Tocris, Bristol, United Kingdom) in 0.5% carboxymethyl cellulose followed by 3-weeks chronic hypoxia (9-10% O_2_), as previously described.^1, 2^ Following completion of hypoxia treatment, the rats were housed under normoxic condition for additional 4 weeks.

**Left main pulmonary artery banding (LPAB)**

Rats were anesthetized in an induction chamber using isoflurane inhalation (5% isoflurane, 2 L/min O_2_). Immediately post-induction, tracheal intubation was performed using a 14 G blunt angiocath that was used to ventilate the rats and maintain anesthesia throughout the entire procedure (tidal volume: 1.5 mL; respiratory rate: 100 breaths/min; 2% isoflurane, 2 L/min O_2_). After confirming surgical plane anesthesia and successful mechanical ventilation, a left thoracotomy was performed in the third intercostal space followed by the section of the intercostal muscles. The pleura was opened and the left lung was mobilized. The left hilum was located, and a 4-0 silk thread was positioned under the left pulmonary artery. The suture was tied tightly around a 25-gauge needle (outer diameter, 0.51 mm) that was placed alongside the left pulmonary artery. Then, the needle was rapidly removed in order to produce a fixed constricted opening in the lumen equal to the diameter of the needle. Subsequently, the chest was closed with 2-0 polydioxanone loop for intercostal spaces. Muscle and skin layers were closed with 4-0 polypropylene. Rats were disconnected from ventilator. Upon confirmation of spontaneous respiration, the animal will be extubated. Buprenorphine and carprofen injections and topical bupivacaine were used for pain management. Topical bupivacaine was applied immediately after wound closure and twice daily for one day post-surgery. Buprenorphine (s.c., 0.03mg/kg) was administered 1 hour prior to surgery and twice daily for two days post-surgery. In addition, carprofen (s.c., 2.5mg/kg) was administered 1 hour prior to surgery and once daily for two days post-surgery.

**Measurement of RVSP and RV hypertrophy**

RVSP was measured using high-fidelity pressure catheters (Transonic-Scisense Inc., ON, Canada). For RV catheterization at end study, rats were anaesthetized by an intraperitoneal injection of xylazine (7 mg/kg) and ketamine (35 mg/kg). The pressure catheter was inserted into the right jugular vein and advanced through the superior vena cava and right atrium into the RV. Hemodynamic parameters were recorded and analyzed using the LabScribe3 software (iWorx, Dover, NH, USA). At end study, after data acquisition, animals were euthanized by exsanguination under anaesthesia. The heart was excised, and the ventricles were dissected from the atria, the aorta and the pulmonary trunk. The RV and left ventricle (LV) and septum (S) were separated, and RV hypertrophy was calculated by measuring the ratio of RV weight to LV plus septum weight (RV/LV+S, Fulton index). The operators acquiring the RVSP and RV hypertrophy data were blinded to the treatment allocation.

**Lung histological measurements**

The left lobe of the lung was inflated via the trachea with 50:50 OCT/saline solution (Tissue-Tek OCT; Qiagen, Mississauga, ON, Canada) and then removed. The left lobe was then cut into thick cross sections and fixed in 4% paraformaldehyde (PFA) for 24 h, rinsed and washed in PBS for 8 hr and stored in 70% ethanol until the day of paraffin embedding. Tissue blocks were sectioned (5μm thickness) with a microtome (Leica Microsystems, Concord, ON, Canada), placed onto poly‑L‑lysine-coated slides, dried at 37°C for 16 hours and then dewaxed and rehydrated through graded alcohols. For microscopy and quantitative morphometry of the lung, hematoxylin and eosin (H&E) staining was performed with standard protocols. Images were acquired by Panoramic DESK (3DHISTECH, Hungary) scanscope using Panoramic Scanner and analyzed using Panoramic Viewer (3DHISTECH, Hungary). Ten random high-power fields (100X magnification) for each rat were analyzed for media wall thickness, total vessel count and vascular occlusion. Media wall thickness as percent of external diameter was estimated as described previously.^2^ Percent Medial wall thickness = ((distance between the internal and external lamina × 2)/external diameter) × 100. For total vessel count, all the vessels (<100μm) were counted from the 10 random fields. The numbers of distal arterioles (<100 μm) with grade-1 (30-80% occlusion) and grade-2 (>80% occlusion) were quantified from the random fields.

**Cleaved caspase-3 and von Wollebrand Factor immunohistochemistry**

PFA fixed and paraffin embedded tissue were sectioned (5μm thickness) with a microtome (Leica Microsystems, Concord, ON, Canada), placed onto poly‑L‑lysine-coated slides, dried at 37°C for 16 hours and then dewaxed and rehydrated through graded alcohols. Antigen retrieval was performed using Citric Acid Based Antigen Unmasking Solution (Vector Labs, Cat# H3300) according to manufacturer’s protocol. Immunohistochemistry was performed using Rabbit specific HRP/DAB (ABC) Detection IHC Kit (Abcam, Cat# ab64261) according to manufacturer’s protocol. Sequential sections were used for cleaved caspase-3 (Cell Signalling Technologies, Cat# 9661S) and Von Willebrand Factor (vWF, Abcam, Cat# ab6994) immunohistochemistry. The primary antibodies (cleaved caspase-3 at 1:40 and vWF at 1:400) were diluted in 1% BSA in PBS and each section was incubated overnight at 4 °C with 80 μL diluted antibody. Images were acquired by Panoramic DESK (3DHISTECH, Hungary) scanscope using Panoramic Scanner and analyzed using Panoramic Viewer (3DHISTECH, Hungary). Ten random high-power fields (100X magnification) for each rat were analyzed for cleaved caspase-3 positive intimal cells as described previously.^3^

**Immunofluorescence microscopy**

PFA fixed and paraffin embedded tissue were sectioned (5μm thickness) with a microtome (Leica Microsystems, Concord, ON, Canada), placed onto poly‑L‑lysine-coated slides, dried at 37°C for 16 hours and then dewaxed and rehydrated through graded alcohols. Slides were immersed in ddH_2_O for 1 hr before proceeding to the autofluorescence quenching treatment using 0.1 M Glycine in PBS buffer (without Calcium nor Magnesium) for 10 min on an orbital shaker. Slides were then washed for 3 times with PBS (3 min each). After removal of paraffin and microwave treatment in basic antigen retrieval buffer (R&D systems; Cat# CTS013), sections were blocked with 5% BSA (Wisent; Cat# 800-095-EG) in wash buffer containing PBS with 0.1% Tween-20 (Sigma-Aldrich, ON, Canada). Streptavidin/biotin blocking (Vector Laboratories; Cat# SP2002) was performed with minor modification in the incubation time. Polyclonal Rabbit anti-Ki67 (Abcam; Cat# ab15580,) and Biotinylated anti-CD144 (R&D systems; Cat# BAF1002) were incubated at 1:100 dilution and 1:20 dilution overnight at 4°C in a humidified chamber. Binding of the anti-Ki67 or anti-CD144 was detected using Donkey anti-Rabbit secondary antibody conjugated with Alexa Fluor Plus 488 (Invitrogen; Cat# A32790, 1:300 dilution) and streptavidin conjugated with Alexa Fluor 594 (Invitrogen; Cat# S11227, 1:300 dilution), respectively. Mounting with counterstain reagent DAPI, Vectorshield^®^ PLUS mounting medium (Vector Laboratories; Cat# H1900), was applied to all sections. Sections were imaged using an inverted confocal microscope (Zeiss, Cat# LSM900) and image analysis was performed using Zen lite Microscopy Software (Zeiss).

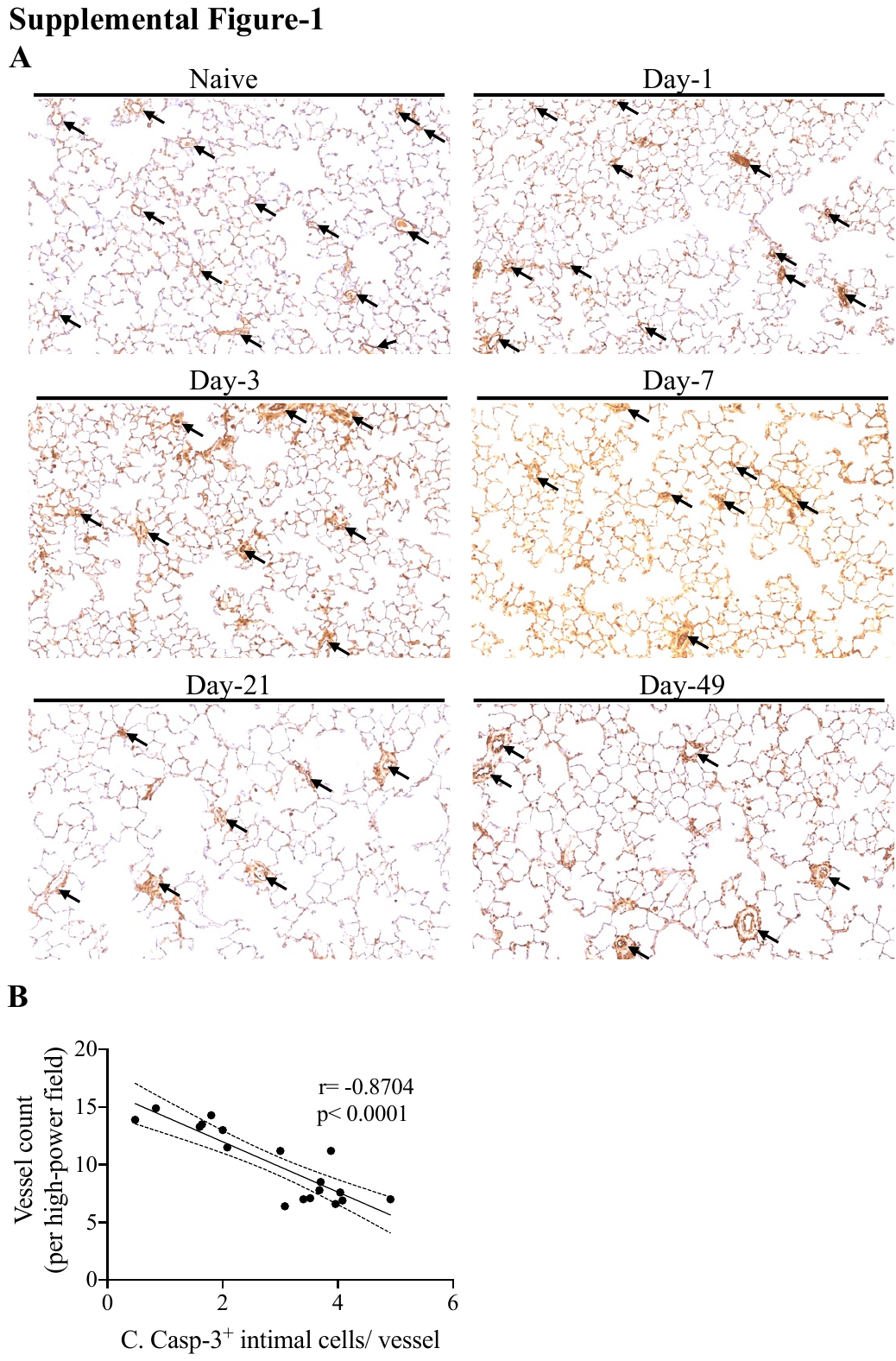

**Supplemental Figure 1: A)** Representative micrograph images demonstrating vessel count at different time-points post-SU in rat SUHx model. **B)** Dot plot showing correlation between cleaved caspase-3 positive intimal cell count per vessel and vessel count per high-power field in the lungs of rats subjected to SUHx model.

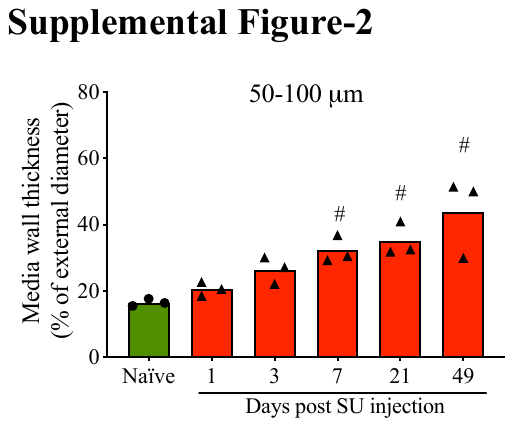

**Supplemental Figure 2:** Media wall thickness of 50-100 μm diameter vessels of male SD rats subjected to SUHx model at different time-points post-SU, N=3 per group, bar graph showing mean, ^#^p<0.05 vs naïve rats

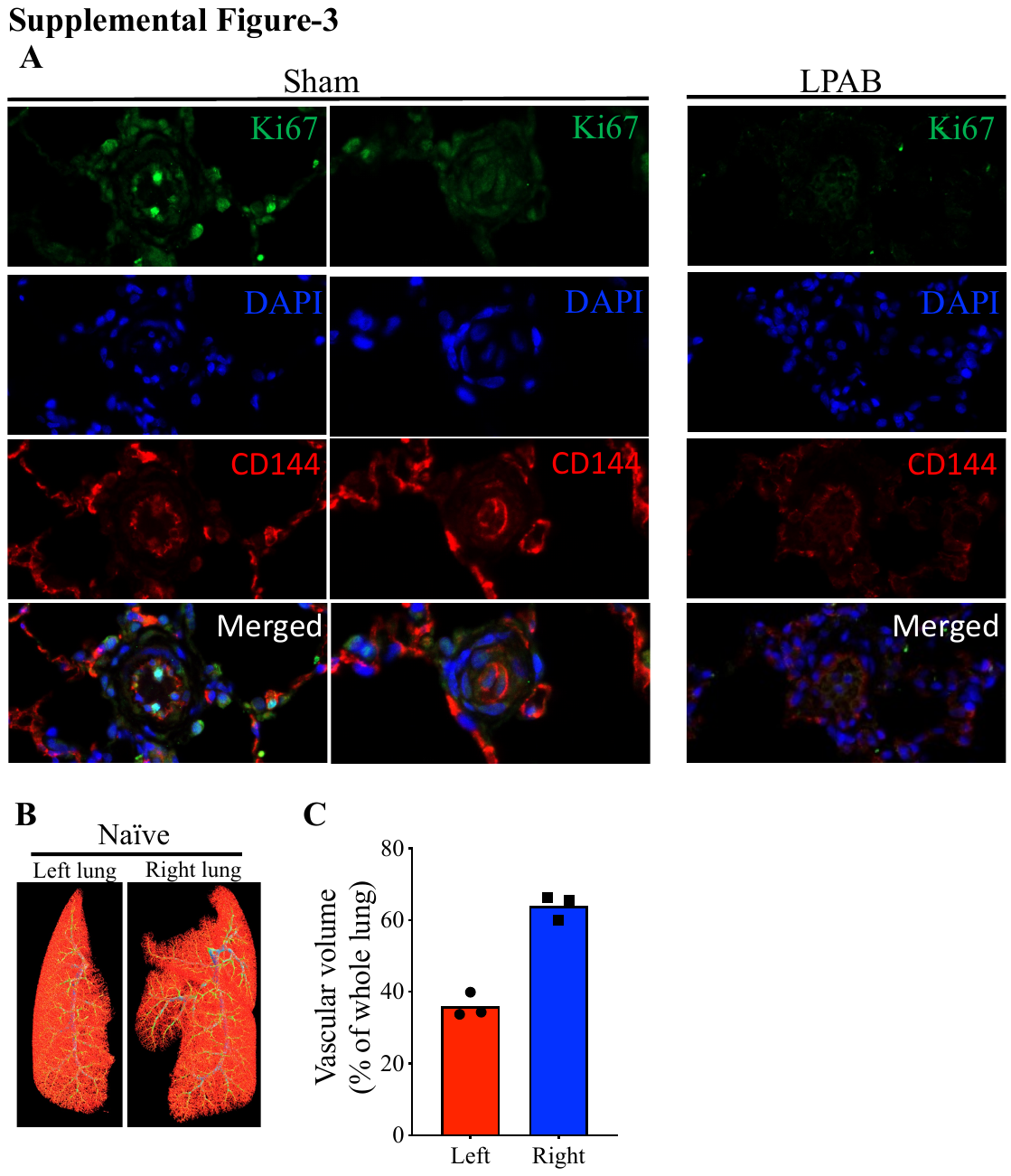

**Figure 3: A)** Representative images demonstrating Ki67 (proliferating cells) and CD144 (endothelial cell marker) staining in pulmonary vasculature of SUHx rats subjected to LPAB or sham procedure. LPAB or sham procedure was performed at 5 weeks post-SU and samples were collected at 8 weeks post-SU **B)** Representative microCT images and **C)** bar graph demonstrating differences in vascular volume of left and right lung of naïve rats. N=3 per group, bar graph showing mean.

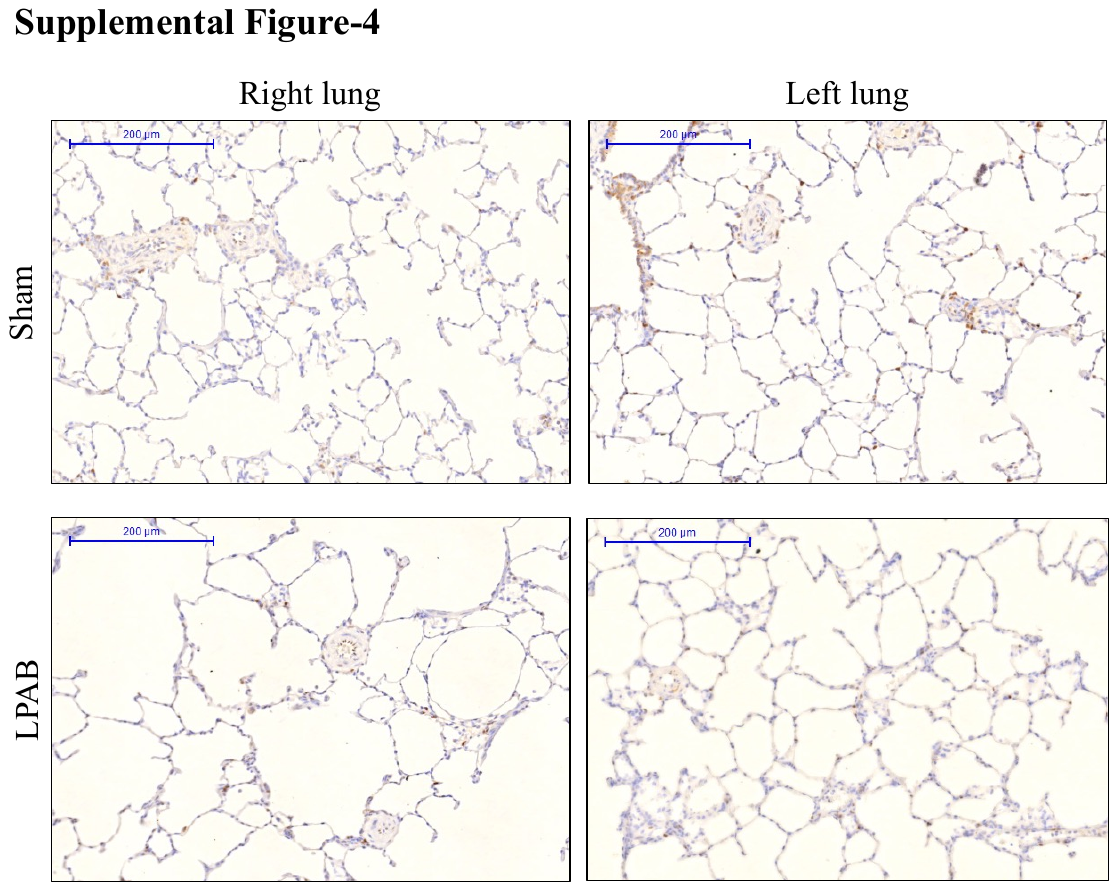

**Figure 4:** Representative images demonstrating cleaved caspase-3 staining in pulmonary vasculature of left and right lung of SUHx rats subjected to LPAB or sham procedure. LPAB or sham procedure was performed at 5 weeks post-SU and samples were collected at 8 weeks post-SU.
